# The genetic basis and evolution of red blood cell sickling in deer

**DOI:** 10.1101/155903

**Authors:** Alexander Esin, L. Therese Bergendahl, Vincent Savolainen, Joseph A. Marsh, Tobias Warnecke

## Abstract

Crescent-shaped red blood cells, the hallmark of sickle cell disease, present a striking departure from the biconcave disc shape normally found in mammals. Characterized by increased mechanical fragility, sickled cells promote haemolytic anaemia and vaso-occlusions and contribute directly to disease in humans. Remarkably, a similar sickle-shaped morphology has been observed in erythrocytes from several deer species, without pathological consequences. The genetic basis of erythrocyte sickling in deer, however, remains unknown, limiting the utility of deer as comparative models for sickling. Here, we determine the sequences of human β-globin orthologs in 15 deer species and identify a set of co-evolving, structurally related residues that distinguish sickling from non-sickling deer. Protein structural modelling indicates a sickling mechanism distinct from human sickle cell disease, coordinated by a derived valine (E22V) in the second alpha helix of the β-globin protein. The evolutionary history of deer β-globins is characterized by incomplete lineage sorting, episodes of gene conversion between adult and foetal β-globin paralogs, and the presence of a trans-species polymorphism that is best explained by long-term balancing selection, suggesting that sickling in deer is adaptive. Our results reveal structural and evolutionary parallels and differences in erythrocyte sickling between human and deer, with implications for understanding the ecological regimes and molecular architectures that favour the evolution of this dramatic change in erythrocyte shape.

## Introduction

Human sickling is caused by a single amino acid change (E6V) in the adult β-globin (HBB) protein (*1*). Upon deoxygenation, steric changes in the haemoglobin tetramer enable an interaction between 6V and a hydrophobic acceptor pocket (known as the EF pocket) on the β-surface of a second tetramer (*2*, *3*). This interaction promotes polymerization of mutant haemoglobin (HbS) molecules, which ultimately coerces red blood cells into the characteristic sickle shape. Heterozygote carriers of the HbS allele are typically asymptomatic (*4*) whereas HbS homozygosity has severe pathological consequences and is linked to shortened lifespan (*5*). Despite this, the HbS allele has been maintained in sub-Saharan Africa by balancing selection because it confers – by incompletely understood means – a degree of protection against the effects of *Plasmodium* infection and malaria (*6*).

Sickling red blood cells were first described in 1840 – seventy years prior to their discovery in humans (*7*) – when Gulliver (*8*) reported unusual erythrocyte shapes in blood from white-tailed deer (*Odocoileus virginianus*). Subsequent research spanning more than a century revealed that sickling, at least as an *in vitro* phenotype, is widespread amongst deer species worldwide (*8*-*11*) (Fig. 1, Table S1). It is not, however, universal: red blood cells from reindeer (*Rangifer tarandus*) and European elk (*Alces alces*, known as moose in North America) do not sickle; neither do erythrocytes from most North American wapiti [*Cervus canadensis*, 25 out of 27 sampled in (*12*); 5 out of 5 sampled in (*11*)]. Below, we will use the term moose for *A. alces* to avoid confusion, as wapiti are also commonly referred to as elk in North America.

**Fig. 1.**
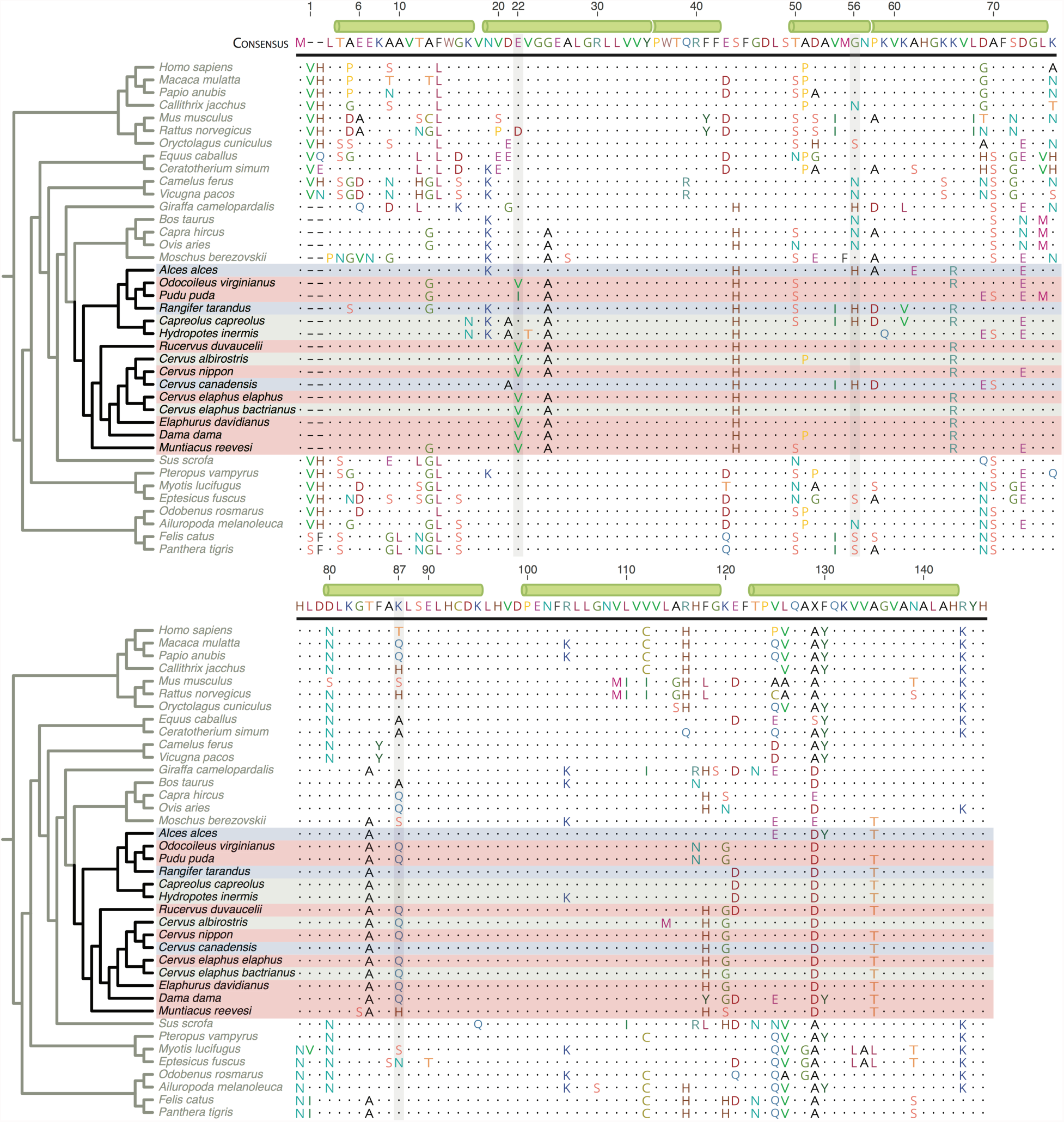
Mammalian adult β-globin peptide sequences in phylogenetic context. To facilitate comparisons with prior classic literature, residues here and in the main text are numbered according to human HBB, skipping the leading methionine. Dots represent residues identical to the consensus sequences, defined by the most common amino acid (X indicates a tie). Key residues discussed in the text are highlighted. β-globin sequences from deer are coloured according to documented sickling state: red = sickling, blue = non-sickling, grey = indeterminate (Table S1). Green cylinders highlight the position of α-helices in the secondary structure of human HBB.

Sickling deer erythrocytes are similar to human HbS cells with regard to their gross morphology and the tubular ultrastructure of haemoglobin polymers (*13*-*16*). Moreover, as in humans, sickling is reversible through modulation of oxygen supply or pH (*9*,*17*). As in humans, deer sickling is mediated by specific β-globin alleles (*18*, *19*), with both sickling and non-sickling alleles segregating in wild populations of white-tailed deer (*20*). As in humans, foetal haemoglobin does not sickle under the same conditions (*19*) and α-globin – two copies of which join two β-globin proteins to form the haemoglobin tetramer - is not directly implicated in sickling etiology (*18*, *21*).

At the same time, a suit of striking differences emerged: whereas human sickling occurs when oxygen tension is low, deer erythrocytes sickle under high pO_2_ and at alkaline pH (*17*). Unlike in humans, the sickling allele (previously labelled β^III^) in white-tailed deer – the only species where allelic diversity has been explored systematically – is the major allele, with a large fraction of individuals (≥60%) homozygous for β^III^ (*20*, *22*). Finally, partial peptide digests suggested that sickling white-tailed deer did not carry the E6V mutation that causes sickling in humans (*22*), leaving the genetic basis of sickling in deer unresolved.

In addition, whereas sickling in humans is unequivocally harmful, it is not known whether sickling in deer is similarly costly, inconsequential or adaptive. Sickling is evident *in vitro* in most deer species, has been induced *in vivo* in sika deer (*Cervus nippon*) by intravenous administration of sodium bicarbonate (*23*) and can be triggered by exercise regimes that lead to transient respiratory alkalosis (*24*).

However, whether sickling occurs intravascularly at appreciable frequency under physiological conditions remains uncertain (*23*). Animals homozygous for the sickling allele do not display aberrant haematological values or other pathological traits (*25*), in line with findings that – unlike human sickle cells – sickled deer erythrocytes do not exhibit increased mechanical fragility *in vitro* (*17*,*18*).

## Results and Discussion

### The molecular basis of sickling in deer is distinct from the human disease state

To dissect the molecular basis of sickling in deer and elucidate its evolutionary history and potential adaptive significance, we used a combination of whole-genome sequencing, locus-specific assembly and targeted amplification to determine the sequences of β-globin genes in 15 deer species, including sickling and non-sickling taxa (Fig. 1, Table S1). Globin genes in mammals are located in paralog clusters, which – despite a broadly conserved architecture – constitute hotbeds of pseudogenization, gene duplication, conversion, and loss (*26*, *27*). In ruminants in particular, globin cluster evolution is highly dynamic. The entire β-globin cluster is triplicated in goat (*Capra hircus*) (*28*) and duplicated in cattle (*Bos taurus*) (*29*), where the two copies of the ancestral β-globin gene sub-functionalized to become specifically expressed in adult (HBB_A_) and foetal (HBB_F_) blood. Consistent with this duplication event pre-dating the Bovidae-Cervidae split, primers designed to amplify HBB_A_ in both these families frequently co-amplified HBB_F_ (see Materials and Methods, Fig. S1c,d). In the first instance, we assigned foetal and adult status based on residues specifically shared with either HBB_A_ or HBB_F_ in cattle, resulting in independent clustering of putative HBB_A_ and HBB_F_ genes on an HBB_A/F_ gene tree (Fig. S2). To confirm these assignments, we sequenced mRNA from the blood (red cell component) of an adult Père David’s deer (*Elaphurus davidianus*) and assembled the erythrocyte transcriptome *de novo* (see Materials and Methods). We identified a highly abundant β-globin transcript (>200,000 transcripts/million, Fig. S3a) corresponding precisely to the putative adult β-globin gene amplified from genomic DNA of the same individual (Fig. S3b). Reads that uniquely matched the putative HBB_A_ gene were >2000-fold more abundant than reads uniquely matching the putative HBB_F_ gene (see Materials and Methods), which is expressed at low levels, is the case in human adults (*30*). Our assignments are further consistent with peptide sequences for white-tailed deer (*22*), fallow deer (*Dama dama*) (*31*) and reindeer (*32*) that were previously obtained from the blood of adult individuals. We then considered deer HBB_A_ orthologs in a wider mammalian context, restricting analysis to species with high-confidence HBB assignments (see Materials and Methods). Treating wapiti as non-sickling, and four species as indeterminate (no or insufficient phenotyping of sickling; Table S1), we find three residues (Fig. 1) that discriminate sickling from non-sickling species: 22 (non-sickling: E, sickling: V/I), 56 (*n-s*: H, *s*: G), and 87 (*n-s*: K, *s*: Q/H). The change at residue 22, from an ancestral glutamic acid to a derived valine (isoleucine in *Pudu puda*) is reminiscent of the human HbS mutation and occurs at a site that is otherwise highly conserved throughout mammalian evolution. The only other amino acid state at residue 22 in our alignment is a biochemically conservative change to aspartic acid (D) in the brown rat (*Rattus norvegicus*).

### Structural modelling supports an interaction between 22V and the EF pocket

To understand how sickling-associated amino acids promote polymerization, we examined these residues in their protein structural context. Residue 22 lies on the surface of the haemoglobin tetramer, at the start of the second alpha helix (Fig. 2a). Close to residue 22 are residue 56 and two other residues that differ between non-sickling reindeer and moose (but not wapiti) and established sickling species: 19 (*n-s*: K, *s*: N) and 120 (*n-s*: K, *s*: G/S). Together these residues form part of a surface of increased hydrophobicity in sickling species (Fig. 2b). Distal to this surface, residue 87 is situated at the perimeter of the EF pocket, which in humans interacts with 6V to laterally link two β-globin molecules in different haemoglobin tetramers and stabilize the parallel strand architecture of the HbS fibre (*2*, *3*, *33*, *34*). Mutation of residue 87 in humans can have marked effects on sickling dynamics (*35*). For example, erythrocytes derived from HbS/Hb Quebec-Chori (T87I) compound heterozygotes sickle like HbS homozygotes (*36*) while Hb D-Ibadan (T87K) inhibits sickling (*37*).

**Fig. 2.**
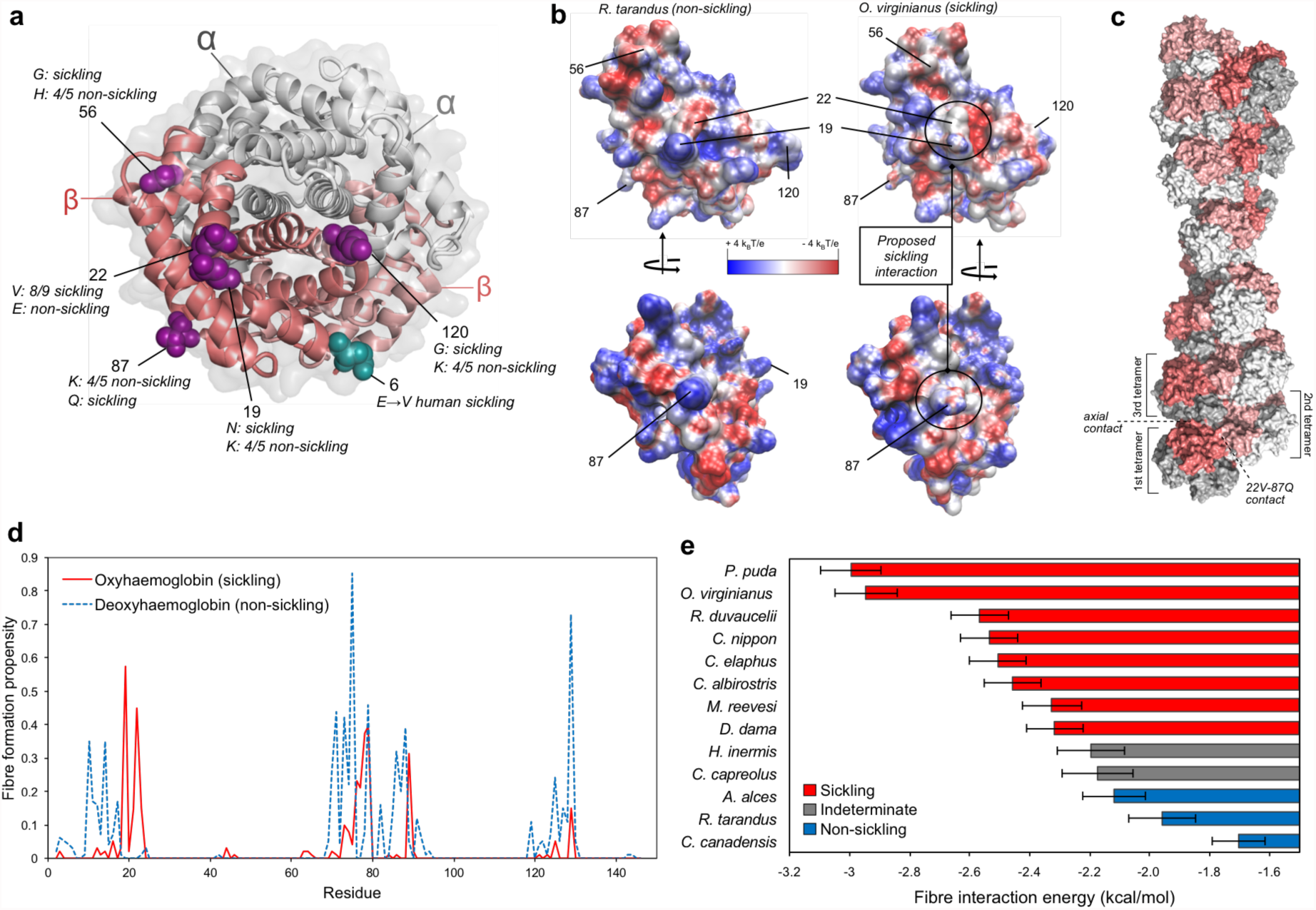
Structural basis for sicking of deer haemoglobin. **a**, Structure of oxyhaemoglobin (PDB ID: 1HHO), with the key residues associated with sickling highlighted in one of the β-globin chains. **b**, Comparison of the electrostatic surfaces of oxy β-globin from a non-sickling (*R. tarandus*) and sickling (*O. virginianus*) species. **c**, Example of a haemoglobin fibre formed via directed docking between residues 22V and 87Q of *O. virginianus* oxy β-globin. **d**, Fibre formation propensity derived from docking simulations centred at a given focal residue in *O. virginianus* oxy and deoxy β-globin. These values represent the fraction of docking models that result in HbS-like haemoglobin fibre structures. **e**, Fibre interaction energy for different deer species, determined by mutating the 270 22V-87Q docking models compatible with fibre formation and calculating the energy of the interaction. Error bars represent standard error of the mean.

Given the similarity between the human E6V mutation and E22V in sickling deer, we hypothesized that sickling occurs through an interaction in *trans* between residue 22 and the EF pocket. To test whether such an interaction is compatible with fibre formation, we carried out directed docking simulations centred on these two residues using a homology model of oxy β-globin from white-tailed deer (see Materials and Methods). We then used the homodimeric interactions from docking to build polymeric haemoglobin structures, analogous to how the 6V-EF interaction leads to extended fibres in HbS homozygotes. Strikingly, nearly half of our docking models resulted in HbS-like straight, parallel strand fibres (Fig. 2c). In contrast, when we performed similar docking simulations centred on residues other than 22V, nearly all were incompatible or much less compatible with fibre formation (Fig. 2d). Out of all 145 β-globin residues, only 19N, which forms a contiguous surface with 22V, has a higher propensity to form HbS-like fibres. By contrast, when docking is carried out using the deoxy β-globin structure 22V is incompatible with fibre formation, consistent with the observation that sickling in deer occurs under oxygenated conditions. Importantly, when this methodology is applied to human HbS, we find that 6V has the highest fibre formation propensity out of all residues under deoxy conditions (Fig. S4a), providing validation for the approach.

Next, we used a force field model to compare the energetics of fibre formation across deer species. We find that known non-sickling species and species suspected to be non-sickling based on their β-globin primary sequence (Chinese water deer, roe deer) exhibit energy terms less favourable to fibre formation than sickling species (Fig. 2e). To elucidate the relative contribution of 22V and other residues to fibre formation, we introduced all single amino acid differences found amongst adult deer β-globin individually into a sickling (*O. virginianus*) and non-sickling (*R. tarandus*) background *in silico* and considered the change in fibre interaction energy. Changes at residue 22 have the strongest predicted effect on fibre formation, along with two residues –19 and 21 – in its immediate vicinity (Fig. S4b). Smaller effects of amino acid substitutions at residue 87, as well as residues 117 (N in *P. puda* and *O. virginianus*) and 118 (Y in *D. dama*) hint at species-specific modulation of sickling propensity. Taken together, the results support the formation of HbS-like fibres in sickling deer erythrocytes via surface interactions centred on residues 22V and 87Q in β-globin molecules of different haemoglobin tetramers.

### Evidence for incomplete lineage sorting during the evolution of *HBB_A_*

To shed light on the evolutionary history of sickling and elucidate its potential adaptive significance, we considered sickling and non-sickling genotypes in phylogenetic context. First, we note that the HBB_A_ gene tree and the species tree (derived from 20 mitochondrial and nuclear genes, see Materials and Methods) are highly discordant (Fig. 3a). Notably, non-sickling and sickling genotypes are polyphyletic on the species tree but monophyletic on the HBB_A_ tree where wapiti, an Old World deer, clusters with moose and reindeer, two New World deer. Gene tree-species tree discordance can result from a number of evolutionary processes, including incomplete lineage sorting, gene conversion, introgression, and classic convergent evolution, where point mutations arise and fix independently in different lineages. In our case, the convergent evolution scenario fits the data poorly. Discordant amino acid states are found throughout the HBB_A_ sequence and are not limited to sickling-related residues (see, for example, the tract of amino acids between residue 44 and 66 shared by *R. tarandus* and *C. capreolus* in Fig. 1). Furthermore, in many instances, amino acids shared between phylogenetically distant species are encoded by the same underlying codons. Conspicuously, this includes the case of residue 120 where all three codon positions differ between sickling species (GGT/AGT) and non-sickling relatives (AAG in reindeer, moose, and the non-sickling ancestor; Fig. S5, Supplementary Data File 1). Even if convergence were driven by selection on a narrow adaptive path through genotype space, precise coincidence of mutational paths at multiple non-synonymous and synonymous sites must be considered unlikely. Rather, these patterns are *prima facie* consistent with incomplete lineage sorting.

**Fig. 3.**
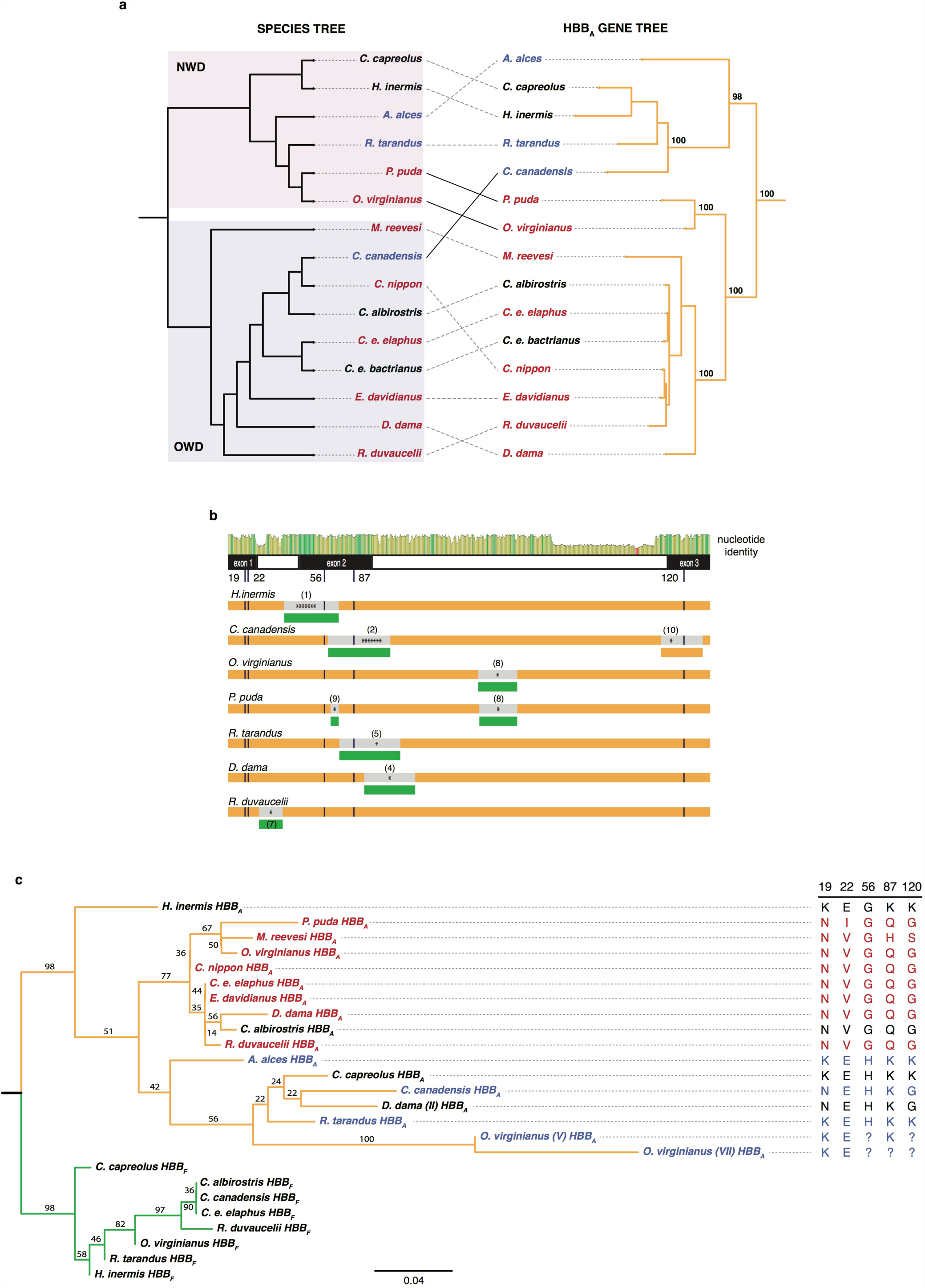
Evidence for incomplete lineage sorting, gene conversion, and a trans-species polymorphism in the evolutionary history of deer HBB_A_. **a,** Discordances between the maximum likelihood HBB_A_ gene tree and the species tree. Topological differences that violate the principal division into New World deer (NWD, Capreolinae) and Old World deer (OWD, Cervinae) are highlighted by solid black lines. Bootstrap values (% out of 1000 bootstrap replicates) are highlighted for salient nodes. **b**, Gene conversion and/or introgression. The top panel illustrates nucleotide identity between HBB_A_ and HBB_F_ orthologs (green: 100%, yellow: 30-100%, red: <30% identity). The low-identity segment towards the end of intron 2 marks a repeat elements present in all adult but absent from all foetal sequences. Below, predicted recombination events affecting HBB_A_ genes (orange), with either an adult ortholog (orange) or a foetal HBB_F_ paralog (green) as the predicted source, suggestive of introgression or gene conversion, respectively. The number of asterisks indicates how many detection methods (out of a maximum of seven) predicted a given event (see Materials and Methods). Details for individual events (numbered in parentheses) are given in Fig. S6a. **c,** Maximum likelihood tree of adult (orange) and foetal (green) β-globin proteins. Alternate non-sickling *D. dama* (II) and *O. virginianus* (V, VII) alleles group with non-sickling species (coloured as in Fig. 1). Amino acid identity at key sites is shown on the right. ?: amino acid unresolved in primary source.

### Gene conversion affects HBB_A_ evolution but does explain the phyletic pattern not of sickling

To shore up this conclusion and rule out alternative evolutionary scenarios, we next asked whether identical genotypes, rather than originating from *de novo* mutations, might have been independently reconstituted from genetic diversity already present in other species (via introgression) or in other parts of the genome (via gene conversion). To evaluate the likelihood of introgression and particularly gene conversion, which has been attributed a prominent role in the evolution of mammalian globin genes (*26*), we first searched for evidence of recombination in an alignment of deer HBB_A_ and HBB_F_ genes. HBB_F_, the most recently diverged paralog of HBB_A_ and itself refractory to sickling (*19*), is the prime candidate to donate non-sickling residues to HBB_A_ in a conversion event. Using a combination of phylogeny-based and probabilistic detection methods and applying permissive criteria that allow inference of shorter recombinant tracts (see Materials and Methods), we identify eight candidate HBB_F_-toHBB_A_ events, two of which, in Chinese water deer and wapiti, are strongly supported by different methods and likely account for hybrid genotypes in exon 2 (Fig. 3b). In addition, we find evidence for recombination in wapiti exon 3. Unlike regions further upstream, the segment affected is 100% identical to other *Cervus spp*., which might be explained by allelic recombination during incomplete lineage sorting (see below). Note, however, that we find no evidence for gene conversion at residue 22 (Fig. 3b, Fig. S6) even when considering poorly supported candidate events. Recombination between HBB_F_ and/or HBB_A_ genes therefore does not explain the re-appearance of 22V in white-tailed deer and pudu (or 22E in wapiti). Consistent with this, removal of putative recombinant regions does not affect the HBB_F_/HBB_A_ gene tree, with wapiti robustly clustered with other non-sickling species whereas white-tailed deer and pudu cluster with Old World sickling species (Fig. S6c). We further screened raw genome sequencing data from white-tailed deer and wapiti for potential donor sequences beyond HBB_F_, such as HBE or pseudogenized HBD sequences, but did not find additional candidate donors. Thus, although gene conversion is a frequent phenomenon in the history of mammalian globins (*26*) and contributes to evolution of HBB loci in deer, it does not by itself explain the recurrence of key sickling/non-sickling residues. Rather, gene conversion introduces additional complexity on a background of incomplete lineage sorting.

### Balancing selection has maintained ancestral variation in HBB_A_

The presence of incomplete lineage sorting and gene conversion confounds straightforward application of rate-based (dN/dS-type) tests for selection, making it harder to establish whether the sickling genotype is simply tolerated or has been under selection. We therefore examined earlier protein-level data on HBB_A_ allelic diversity in extant deer populations. Intriguingly, we find evidence for long-term maintenance of ancestral variation. Two rare non-sickling β-globin alleles in white-tailed deer (previously identified from partial peptide digests) cluster with the non-sickling β-globin of reindeer rather than with the white-tailed deer sickling allele (Fig. 3c), albeit with modest bootstrap support. In addition, protein-level allelic diversity has also been observed in fallow deer, a sickling Old World deer, where the alternate β-chain (*31*) is closely related to the moose sequence but substantially different from the sickling allele that we recovered in our sample (Fig. 3c). Finally, phenotypic heterogeneity in wapiti (*12*) and sika deer (*38*) sickling indicates that rare sickling and non-sickling variants, respectively, also segregate in these two species. Taken together, these findings point to the long-term maintenance of ancestral variation through successive speciation events dating back to the most common ancestor of Old World and New World deer, an estimated ~13.6 million years ago (mya) [CI: 9.84-17.33mya, (*39*)].

Might this polymorphism have been maintained simply by chance or must balancing selection be evoked to account for its survival? We currently lack information on broader patterns of genetic diversity at deer HBB_A_ loci and surrounding regions that would allow us to search for footprints of balancing selection explicitly. However, we can estimate the probability *P* that a trans-species polymorphism has been maintained along two independent lineages by neutral processes alone as

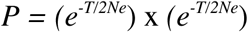

where *T* is the number of generations since the two lineages split and *N_e_* is the effective population size (*40*, *41*). For simplicity, *N_e_* is assumed to be constant over *T* and the same for both lineages. In the absence of reliable species-wide estimates for *N_e_*, we can nonetheless ask what *N_e_* would be required to meet a given threshold probability. Conservatively assuming an average generation time of 1 year (*42*, *43*) and a split time of 7.2mya [the lowest divergence time estimate in the literature (*39*)], *N_e_* would have to be 2,403,419 to reach a threshold probability of 0.05 (1,563,460 for *P*=0.01). Although deer populations can have a large census population sizes, an *N_e_* >2,000,000 for both fallow and white-tailed deer is comfortably outside what we would expect for large-bodied mammals, >4-fold higher than estimates for wild mice (*44*) and >2-fold higher even than estimates for African populations of *Drosophila melanogaster* (*45*). Consequently, we posit that the HBB_A_ trans-species polymorphism is inconsistent with neutral evolution and instead reflects the action of balancing selection.

Balanced polymorphisms shared between human and chimp are principally related to immune function and parasite pressure (*46*, *47*), so it is tempting to speculate that similar selection pressures might operate in deer, which host a number of intra-erythrocytic parasites including *Babesia* (*48*) and *Plasmodium* (*49*). Unlike in humans, however, the maintenance of two distinct alleles cannot be attributed to heterozygote advantage: Sickling homozygotes are the norm in white-tailed deer (*20*, *22*, *50*) and not associated with an outward clinical phenotype, indicating that the cost of bearing two sickling alleles must be comparatively low. Why, then, is the non-sickling allele being maintained? One possibility is that the sickling allele is cost-free most of the time, as previously suggested (*24*), but carries a burden in a particular environment, so that the sickling or non-sickling allele might be favoured in different sub-populations depending on local ecology. In this scenario, allelic diversity might be maintained by migration-selection balance. However, what ecological features might define such environments remains an open question. Considering only species for which phenotypic sickling information is available, there is a marked geographic asymmetry in sickling status, where non-sickling species are restricted to arctic and subarctic (elk, reindeer) or mountainous (wapiti) habitat. This might indicate that the sickling allele loses its adaptive value in colder climates (perhaps linked to the lower prevalence of blood-born parasites). However, based on genotype, we would also predict Chinese water deer and roe deer to be amongst the non-sicklers yet these species are widespread in temperate regions and their ranges overlap extensively with sickling species, challenging the hypothesis that ambient temperature is a primary driver of sickling. Future epidemiological studies coupled to ecological and population genetic investigations will be required to unravel the evolutionary ecology of sickling in deer, establish whether parasites are indeed ecological drivers of between- and within-species differences in HBB_A_ genotype and, ultimately, whether deer might serve as a useful comparative system to elucidate the link between sickling and protection from the effects of *Plasmodium* infection, which remains poorly understood in humans.

## Materials and Methods

### Sample collection and processing

Blood, muscle tissue, and DNA samples were acquired for 15 species of deer from a range of sources (Table S1). The white-tailed deer blood sample was heat-treated on import to the United Kingdom in accordance with import standards for ungulate samples from non-EU countries (IMP/GEN/2010/07). Fresh blood was collected into PAXgene Blood DNA tubes (PreAnalytix) and DNA extracted using the PAXgene Blood DNA kit (PreAnalytix). DNA from previously frozen blood samples was extracted using the QIAamp DNA Blood Mini kit (Qiagen). DNA from tissue samples was extracted with the QIAamp DNA Mini kit (Qiagen) using 25mg of tissue. Total RNA was isolated from an *E. davidianus* blood sample using the PAXgene Blood RNA kit (PreAnalytix) three days after collection into a PAXgene Blood RNA tube (PreAnalytix). All extractions were performed according to manufacturers’ protocols. For each sample, we validated species identity by amplifying and sequencing the cytochrome b (*CytB*) gene. With the exception of *Cervus albirostris*, we successfully amplified *CytB* from all samples using primers MTCB_F/R (Fig. S1) and conditions as described in (*51*). Phusion High-Fidelity PCR Master Mix (ThermoFisher) was used for all amplifications. PCR products were purified using the MinElute PCR Purification Kit (Qiagen) and Sanger-sequenced with the amplification primers. The *CytB* sequences obtained were compared to all available deer *CytB* sequences in the 10kTrees Project (*52*) using the *ape* package (function *dist.dna* with default arguments) in R (*53*). In all cases, the presumed species identity of the sample was confirmed (Table S2).

### Whole genome sequencing

*O. virginianus* genomic DNA was prepared for sequencing using the NEB DNA library prep kit (New England Biolabs) and sequenced on the Illumina HiSeq platform. The resulting 229 million paired-end reads were filtered for adapters and quality using Trimmomatic (*54*) with the following parameters: *ILLUMINACLIP:adapters/TruSeq3-PE-2.fa:2:30:10 LEADING:30 TRAILING:30 SLIDINGWINDOW:4:30 MINLEN:50*. Inspection of the remaining 163.5M read pairs with FastQC (http://www.bioinformatics.babraham.ac.uk/projects/fastqc/) suggested that overrepresented sequences had been successfully removed.

### Mapping and partial assembly of the *O. virginianus* β-globin locus

To seed a local assembly of the *O. virginianus* β-globin locus we first mapped *O. virginianus* trimmed paired-end reads to the duplicated β-globin locus in the hard-masked *B. taurus* genome (UMD 3.1.1; chr15: 48973631-49098735). The β-globin locus is defined here as the region including all *B. taurus* β-globin genes (HBE1, HBE4, HBB, HBE2, HBG), the intervening sequences and 24kb either side of the two outer β-globins (HBE1, HBG). The mapping was performed using bowtie2 (*55*) with default settings and the optional *--no-mixed* and *--no-discordant* parameters. 110 reads mapped without gaps and a maximum of one nucleotide mismatch. These reads, broadly dispersed across the *B. taurus* β-globin locus (Fig. S7), were used as seeds for local assembly using a customised aTRAM (*56*) pipeline (see below). Prior to assembly, the remainder of the reads were filtered for repeat sequences by mapping against Cetartiodactyla repeats in Repbase (*57*). The aTRAM.pl wrapper script was modified to accept two new arguments: *max_target_seqs <int>* limited the number of reads found by BLAST from each database shard; *cov_cutoff <int>* passed a minimum coverage cut-off to the underlying Velvet 1.2.10 assembler (*58*). The former modification prevents stalling when the assembly encounters a repeat region, the latter discards low coverage contigs at the assembler level. aTRAM was run with the following arguments: *-kmer 31 -max_target_seqs 2000 -ins_length 270 -exp_coverage 8 -cov_cutoff 2 -iterations 5*. After local assembly on each of the 110 seed reads, the resulting contigs were combined using Minimo (*59*) with a required minimum nucleotide identity of 99%. To focus specifically on assembling the adult βglobin gene, only contigs that mapped against the *B. taurus* adult β-globin gene ±500bp (chr15: 49022500-49025000) were retained and served as seeds for another round of assembly. This procedure was repeated twice. The final 59 contigs were compared to the UMD 3.1.1 genome using BLAT and mapped exclusively to either the adult or foetal *B. taurus* β-globin gene. From the BLAT alignment, we identified short sequences that were perfectly conserved between the assembled deer contigs and the *B. taurus* as well as the sheep (*Ovis aries*) assembly (Oar_v3.1). Initial forward and reverse primers (Ovirg_F1/Ovirg_R1, Fig. S1a) for β-globin amplification were designed from these conserved regions located 270bp upstream (chr15:49022762-49022786) and 170bp downstream (chr15:49024637-49024661) of the *B. taurus* adult β-globin gene, respectively.

### Globin gene amplification and sequencing

Amplification of β-globin from *O. virginianus* using primers Ovirg_F1 and Ovirg_R1 yielded two products of different molecular weights (~2000bp and ~1700bp; Fig. S1c), which were isolated by gel extraction and Sanger-sequenced using the amplification primers. The high molecular weight product had higher nucleotide identity to the adult (93%) than to the foetal (90%) *B. taurus* β-globin coding sequence. Note that the discrepancy in size between the adult and foetal β-globin amplicons derives from the presence of two tandem Bov-tA2 SINEs in intron 2 of the adult β-globin gene in cattle, sheep, and *O. virginianus* and is therefore likely ancestral. We designed a second set of primers to anneal immediately up- and downstream, and in the middle of the adult β-globin gene (Ovirg_F2, Ovirg_R2, Ovirg_Fmid2, Fig. S1a,b). Amplification from DNA extracts of other species with Ovirg_F1/Ovirg_R1 produced mixed results, with some species showing a two-band pattern similar to *O. virginianus*, others only a single band – corresponding to the putative adult β-globin (Fig. S1d). Using these primers, no product could be amplified from *R. tarandus*, *H. inermis*, and *C. capreolus*. We identified a 3bp mismatch to the Ovirg_R1 primer in a partial assembly of *C. capreolus* (Genbank accession: GCA_000751575.1; scaffold: CCMK010226507.1) that is likely at fault. A re-designed reverse primer (Ccap_R1) successfully amplified the adult β-globin gene from the three deer species above as well as *C. canadensis* (Fig. S1). All amplifications were performed using Phusion High-Fidelity PCR Master Mix (ThermoFisher), with primers as listed in Fig. S1a, and 50-100ng of genomic DNA. Annealing temperature and step timing were chosen according to manufacturer guidelines. Amplifications were run for 35 cycles. Gel extractions were performed on samples resolved on 1% agarose gels for 40 minutes at 90V using the MinElute Gel Extraction Kit (Qiagen) and following the manufacturer’s protocol. PCR purifications were performed using the MinElute PCR Purification Kit (Qiagen) following the manufacturer’s protocol. All samples were sequenced using the Sanger method with amplification primers and primer Ovirg_Fmid2.

### Transcriptome sequencing and assembly

RNA was extracted from the red cell component of a blood sample of an adult Père David’s deer using the PAXgene Blood RNA kit (Qiagen). An mRNA library was prepared using a Truseq mRNA library prep kit and sequenced on the MiSeq PE150 platform, yielding 25,406,472 paired-end reads, which were trimmed for adapters and quality-filtered using Trim Galore! (http://www.bioinformatics.babraham.ac.uk/projects/trim_galore/) with a base quality threshold of 30. The trimmed reads were used as input for *de novo* transcriptome assembly with Trinity (*60*) using default parameters. A blastn homology search against these transcripts, using the *O. virginianus* adult β-globin CDS as query, identified a highly homologous transcript (E-value = 0; no gaps; 97.5% sequence identity compared with 92.2% identity to the foetal β-globin). The CDS of this putative β-globin transcript was 100% identical to the sequence amplified from Père David’s deer genomic DNA (Fig. S3b). We used *emsar* (*61*) with default parameters to assess transcript abundances. The three most abundant reconstructed transcripts correspond to full or partial α- and β-globin transcripts, including one transcript, highlighted above, that encompasses the entire adult β-globin CDS. These transcripts are an order of magnitude more abundant than the fourth most abundant (Fig. S3a), in line with the expected predominance of α- and β-globin transcripts in mature adult red blood cells. To investigate whether the foetal β-globin could be detected in the RNA-seq data and because amplification of the foetal β-globin from Père David’s deer genomic DNA was not successful, we mapped reads against the foetal β-globin gene of *C. e. elaphus*, the closest available relative. Given that the CDS of the adult β-globins in these species are 100% identical, we expected that the foetal orthologs would likewise be highly conserved. We therefore removed reads with more than one mismatch and assembled putative transcripts from the remaining 1.3M reads using the Geneious assembler v.10.0.5 (*62*) with default parameters (*fastest* option enabled). We recovered a single contig with high homology to the *C. e. elaphus* foetal β-globin CDS (only a single mismatch across the CDS). We then estimated the relative abundance of adult and the putative foetal transcripts by calculating the proportion of reads that uniquely mapped to either the adult or foetal CDS. 1820532 reads mapped uniquely to the adult sequence whereas 872 mapped uniquely to the foetal CDS, a ratio of 2088:1.

### Structural analysis

Homology models were built for *O. virginianus* and *R. tarandus* β-globin sequences using the MODELLER-9v15 program for comparative protein structure modelling (*63*) using both oxy (1HHO) and deoxy (2HHB) human haemoglobin structures as templates. The structures were used for electrostatic calculations using the Adaptive Poisson-Boltzmann Solver (*64*) plugin in the Visual Molecular Dynamics (VMD) program (*65*). The surface potentials were visualised in VMD with the conventional red and blue colours, for negative and positive potential respectively, set at ±5 kT/e.

### Modelling of haemoglobin fibres

We first used the program HADDOCK (*66*) with the standard protein-protein docking protocol to generate ensembles of docking models of β-globin dimers. In each docking run, a different interacting surface centred around a specific residue was defined on each β-globin chain. All residues within 3Å of the central residue were defined as “active” and were thus constrained to be directly involved in the interface, while other residues within 8Å of the central residue were defined as “passive” and were allowed but not strictly constrained to form a part of the interface. We performed docking runs with the interaction centred between residue 87 and all other residues, generating at least 100 water-refined β-globin dimer models for each (although 600 *O. virginianus* oxy β-globin 22V-87Q models were built for use in the interaction energy calculations). The β-globin dimers were then evaluated for their ability to form HbS-like fibres out of full haemoglobin tetramers. Essentially, the contacts from the β-globin dimer models were used to build a chain of five haemoglobin molecules, in the same way that the contacts between 6V and the EF pocket lead to an extended fibre in HbS. HbS-like fibres were defined as those in which a direct contact was formed between the first and third haemoglobin tetramers in a chain (analogous to the axial contacts in HbS fibres, see Fig. 2c), and in which the chain is approximately linear. This linearity was measured as the distance between the first and third plus the distance between the third and the fifth haemoglobin tetramers, divided by the distance between the first and the fifth. A value of 1 would indicate a perfectly linear fibre, while we considered any chains with a value <1.05 to be approximately linear and HbS-like. Finally, chains containing significant steric clashes between haemoglobin tetramers (defined as >3% of Cα atoms being within 2.8Å of another Cα atom) were excluded. Fibre formation propensity was then defined as the fraction of all docking models that led to HbS-like fibres.

### Interaction energy analysis

Using the 270 22V-87Q models of *O. virginianus* β-globin dimers that can form HbS-like fibres, we used FoldX (*67*) and the ‘RepairPDB’ and ‘BuildModel’ functions to mutate each dimer to the sequences of all other adult deer species. Note that since *C. e. elaphus, C. e. bactrianus* and *E. davidianus* have identical amino acid sequence, only one of these was included here. The energy of the interaction was then calculated using the ‘AnalyseComplex’ function of FoldX, and then averaged over all docking models. The same protocol was then used for the analysis of the effects of individual mutations, using all possible single amino acid substitutions observed in the adult deer sequences, except that the interaction energy was presented as the change with respect to the wild-type sequence.

### Detection of recombination events

We considered two sources of donor sequence for recombination into adult β-globins: adult β-globin orthologs in other deer species and the foetal β-globin paralog within the same genome. *H. inermis* HBB_F_ was omitted from this analysis since the sequence of intron 2 was only partially determined. We used the Recombination Detection Program (RDP v.4.83) (*68*) to test for signals of recombination in an alignment of complete adult and foetal deer β-globin genes that were successfully amplified and sequenced, enabling all subtended detection methods (including primary scans for BootScan and SiScan) except LARD, treating the sequences as linear and listing all detectable events. In humans, conversion tracts of lengths as short as 110bp have been detected in the globin genes (*69*) and tracts as short as 50bp in other gene conversion hotspots (*70*, *71*). Given the presence of multiple regions of 100% nucleotide identity across the alignment of adult and foetal deer β-globins (Fig. 3b), we suspected that equally short conversion tracts might also be present. We therefore lowered window and step sizes for all applicable detection methods in RDP (Fig. S6b) at the cost of a lower signal-to-noise ratio. As the objective is to test whether recombination events could have generated the phyletic distribution of sickling/non-sickling genotypes observed empirically, this is conservative.

### Phylogenetic analyses

Adult deer β-globin coding sequences were aligned with MUSCLE to a set of non-chimeric mammalian β-globin CDSs (*26*). The mammalian phylogeny (Fig. S8) is principally based on the Timetree of Life (*39*) with the order Carnivora regrafted to branch above the root of the Chiroptera and Artiodactyla to match findings in (*72*). The internal topology of Cervidae was taken from the Cetartiodactyla consensus tree of the 10kTrees Project (*52*). *C. canadensis* and *Cervus elaphus bactrianus*, not included in the 10kTrees phylogeny, were added as sister branches to *C. nippon* and *C. e. elaphus*, respectively, following (*73*).

### Data availability

HBB_A_ and HBB_F_ full gene sequences (coding sequence plus intervening introns) have been submitted to GenBank with accession numbers KY800429-KY800452. Père David’s deer RNA sequencing and white-tailed deer whole genome sequencing raw data has been submitted to the European Nucleotide Archive (ENA) with the accession numbers PRJEB20046 and PRJEB20034, respectively.

## Acknowledgments

We thank the Zoological Society of London Whipsnade Zoo (F. Molenaar), Bristol Zoological Society (S. Dow, K. Wyatt), the Royal Zoological Society of Scotland Highland Wildlife Park (J. Morse), the Penn State Deer Research Center (D. Wagner), and the Northeast Wildlife DNA Laboratory (N. Chinnici) for samples, the MRC LMS Genomics Facility for DNA and RNA sequencing, B.N. Sacks, J. Mizzi, and T. Brown for access to Tule elk sequencing data, P.D. Butcher for discussions, and P. Sarkies, A. Brown, and B. Lehner for comments on the manuscript. This work was supported by an Imperial College Interdisciplinary Cross-Campus Studentship to A.E, an MRC Career Development Award (MR/M02122X/1) to J.A.M., a Leverhulme Trust Fellowship to V.S., and MRC core funding and an Imperial College Junior Research Fellowship to T.W.

## Author contributions

A.E. performed laboratory experiments and evolutionary analyses and contributed to experimental design, data analysis and interpretation. L.T.B. and J.A.M. designed and performed structural modelling, and contributed to data analysis and interpretation. V.S. contributed tissue samples. T.W. conceived the study, contributed to experimental design, data analysis, and interpretation and wrote the manuscript with the input from all authors.

## Author information

The authors declare no competing financial interests.

---

**Fig. S1.**
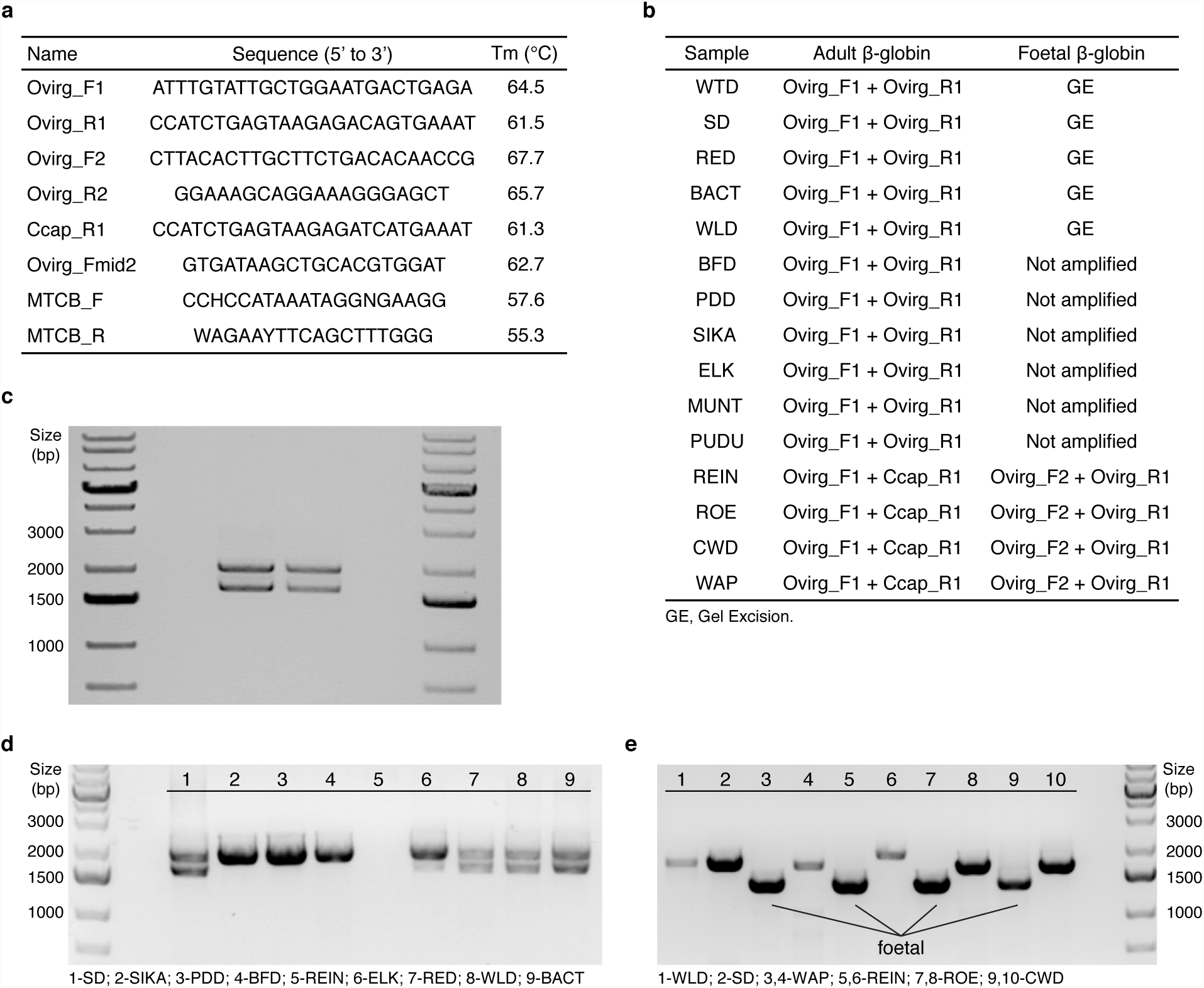
Amplification of deer β-globin genes. **a**, Primers used in this study. **b**, Primer combinations used to amplify adult and foetal β-globin genes in different species. Where possible, gel excision was used to isolate the co-amplified foetal β-globin band. In certain cases, the adult gene could also be selectively amplified using primers Ovirg_F1/Ovirg_R2 (see panel e, lanes 1,2). **c**, Agarose gel showing co-amplification of adult and foetal β-globin genes in white-tailed deer with Ovirg_F1/Ovirg_R1; the two lanes show two different individuals. **d**, Agarose gel showing heterogeneity of amplification products in different deer using Ovirg_F1/Ovirg_R1. **e**, Agarose gel showing selective amplification of adult and foetal β-globin genes. Lanes 1 & 2: adult P-globin using Ovirg_F1/Ovirg_R2; all other lanes with primer combinations described in panel b.

**Fig. S2.**
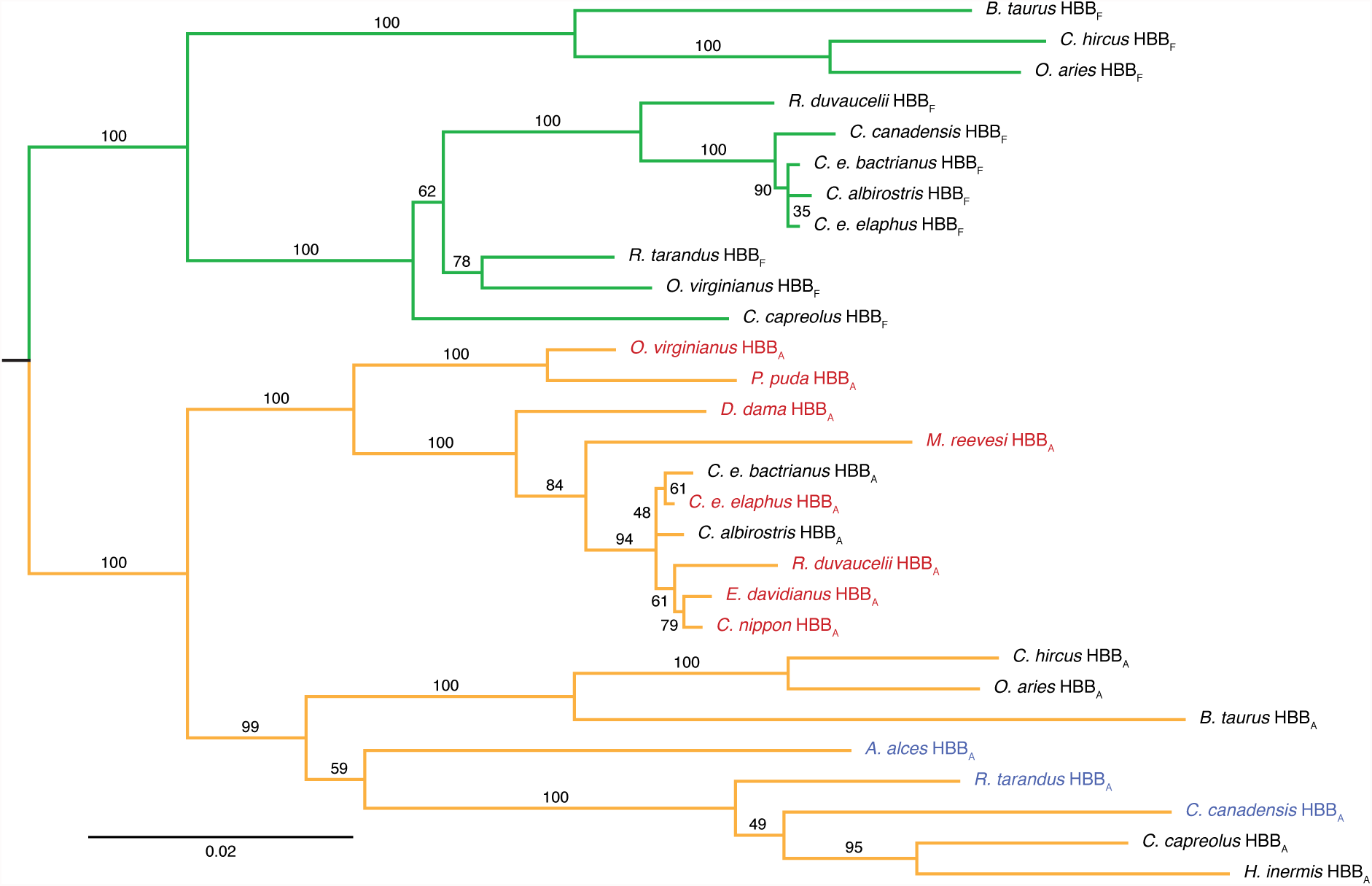
Putative HBB_A_ and HBB_F_ genes cluster separately on an HBB_A_/HBB_F_ gene tree. The tree is a maximum likelihood reconstruction based on a nucleotide alignment of complete exonic and intronic sequences (see Materials and Methods). The sequence of *H. inermis* HBB_F_ was only partially resolved and is hence omitted. Branches are coloured as adult (orange) or foetal (green). Tip labels for HBB_A_ are coloured according to the species’ propensity to sickle where known (red = sickling, blue = non-sickling). Bootstrap values are derived from 100 bootstrap replicates. The scale bar shows the number of nucleotide substitutions per site.

**Fig. S3.**
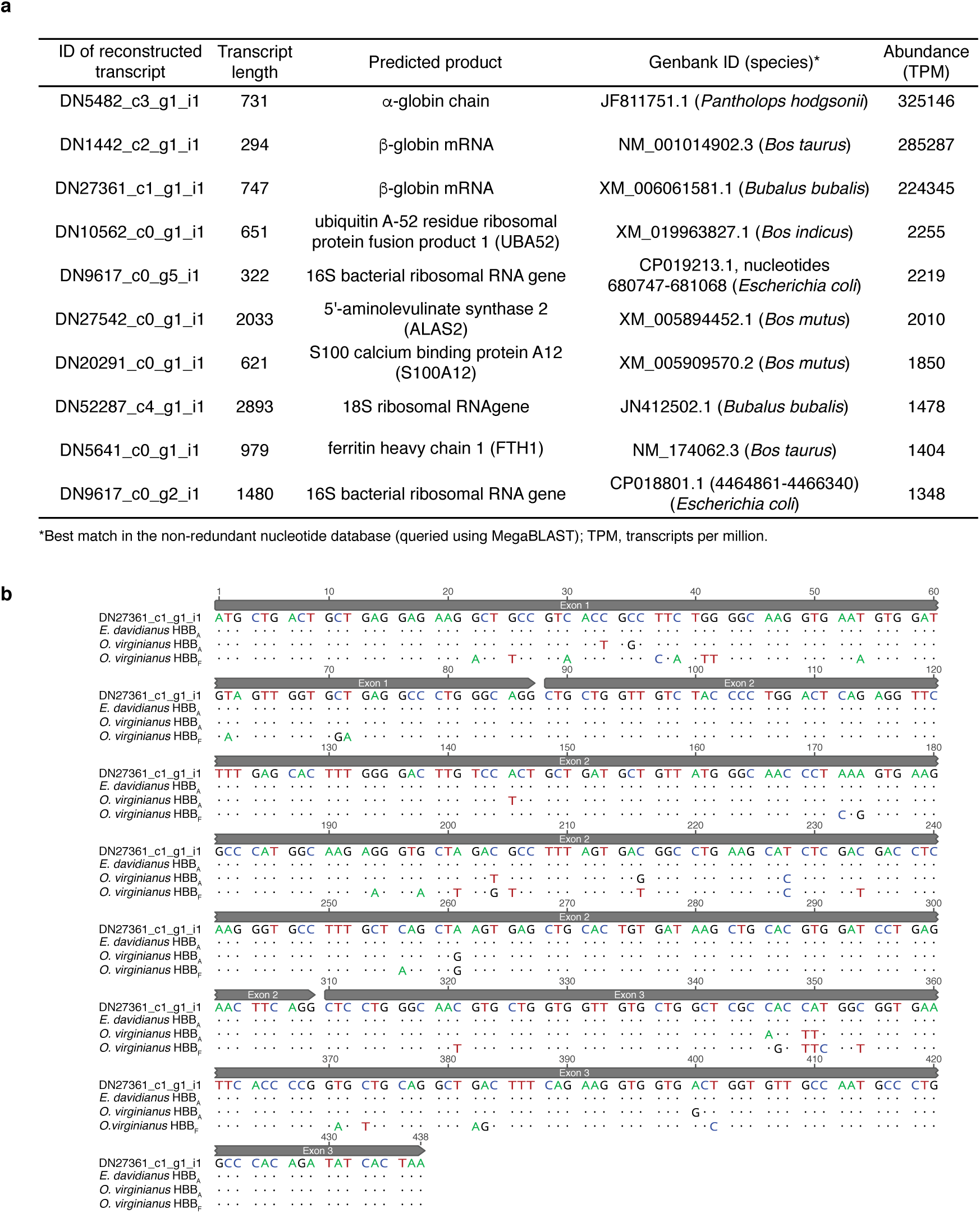
Reconstructing adult β-globin sequence from RNA sequencing data. **a**, the ten most abundant transcripts in the *de novo* assembled *E. davidianus* red blood cell transcriptome. **b**, Nucleotide alignment of the *E. davidianus* β-globin CDS derived from the *de novo* transcriptome assembly, the putative *E. davidianus* adult β-globin CDS derived from amplification, and the foetal and adult β-globin CDSs from *O. virginianus*.

**Fig. S4.**
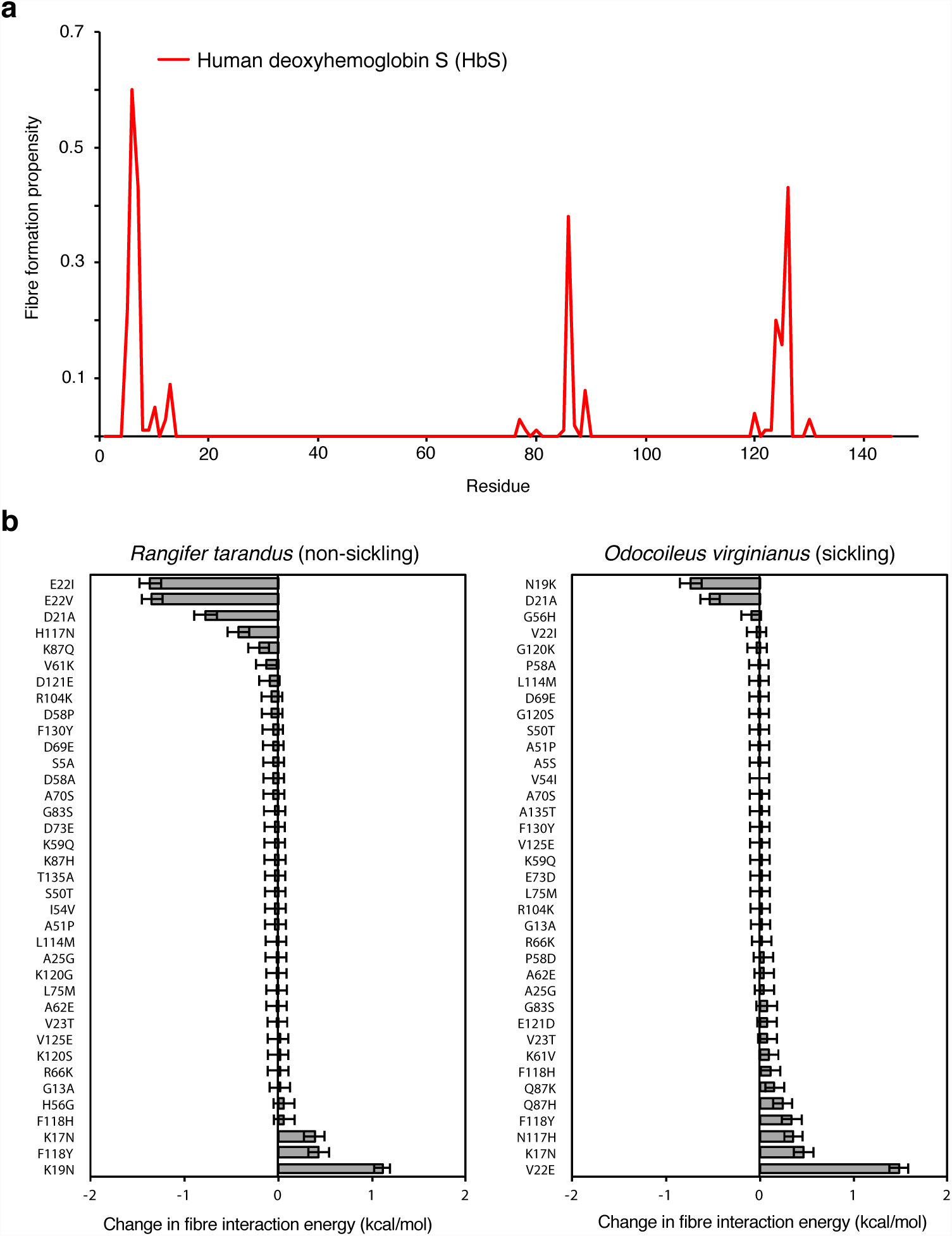
Analysis of sickling propensity in human and deer. **a**, Fibre formation propensity assuming an interaction between the EF pocket and a given focal residue on two different β-globin chains, essentially as in Fig. 2d, but using the structure of human deoxyhaemoglobin S (HbS). Fibre formation propensity represents the fraction of the 100 β-globin dimer models built for each position that can form HbS-like fibres. **b**, Effects on fibre interaction energy of replacing defined single amino acids in the primary sequence of either *R. tarandus* or *O. virginianus*. Negative values indicate stronger interactions and thus an increased likelihood of fibre formation. These values show the mean over all 270 22V-87Q docking models compatible with fibre formation, and error bars represent the standard error of the mean.

**Fig. S5.**
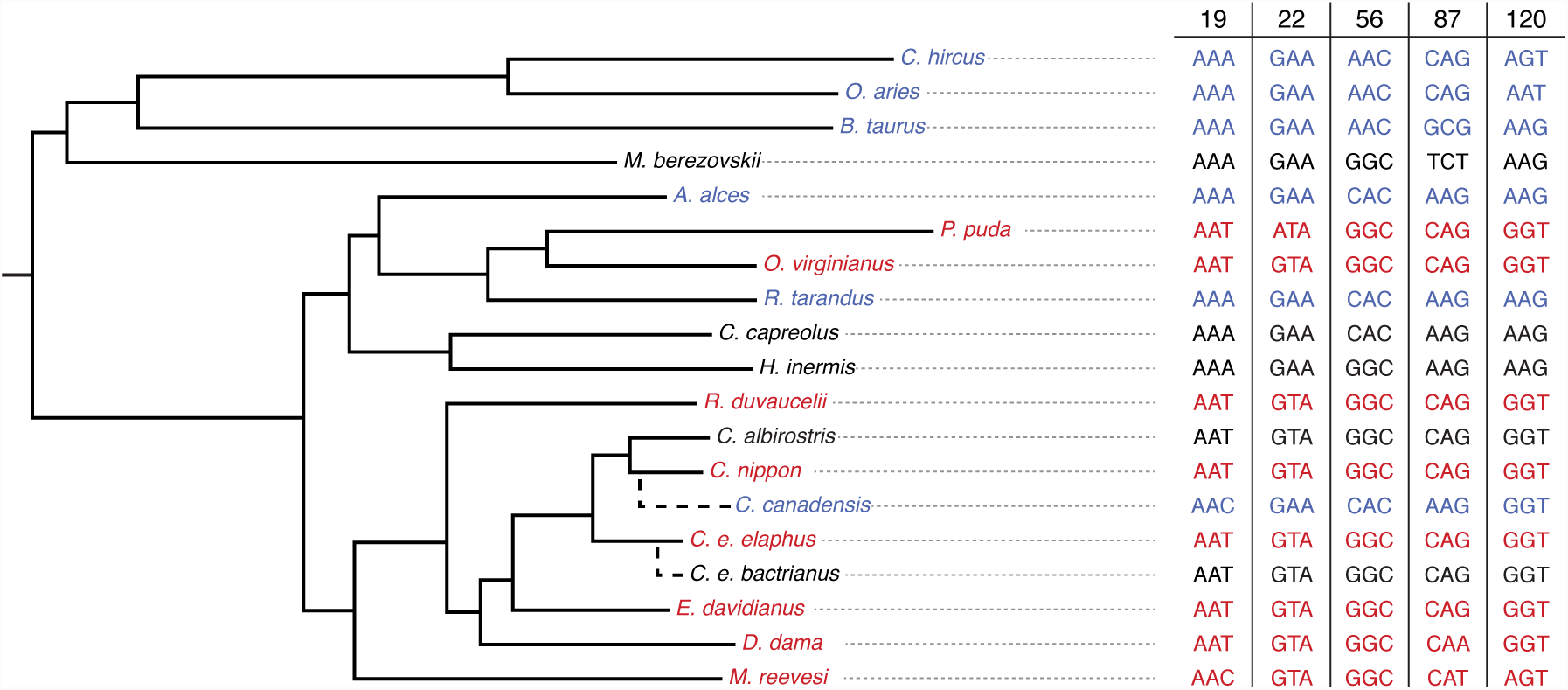
Codons specifying sickling-associated amino acids in HBB_A_ genes of different deer species. Tip labels and associated codons coloured as in Fig. 3a

**Fig. S6.**
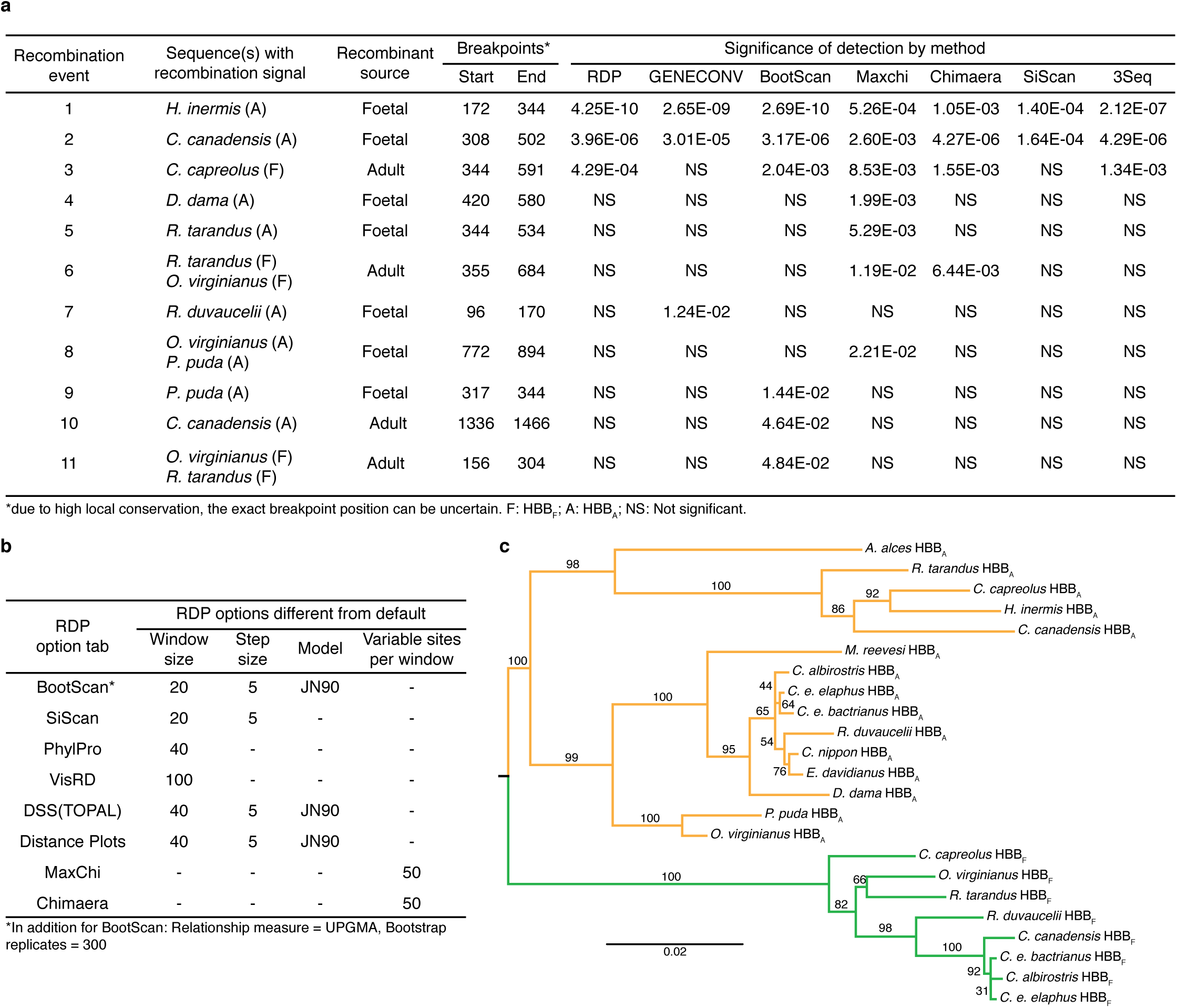
Detection of gene conversion and introgression events in deer β-globin genes. **a**, Recombination events predicted from an alignment of deer HBB_F_ and HBB_A_ genes by different methods. Breakpoint positions are given relative to each focal sequence. Where two sequences are affected (events 6,8,11) positions refer to the top sequence. **b**, Non-default parameters used for detecting recombination events with RDP. **c**, Maximum likelihood tree derived from the alignment of adult (orange) and foetal (green) β-globin genes after predicted recombinant regions have been removed. Branch support values derived from 100 bootstrap replicates are given.

**Fig. S7.**
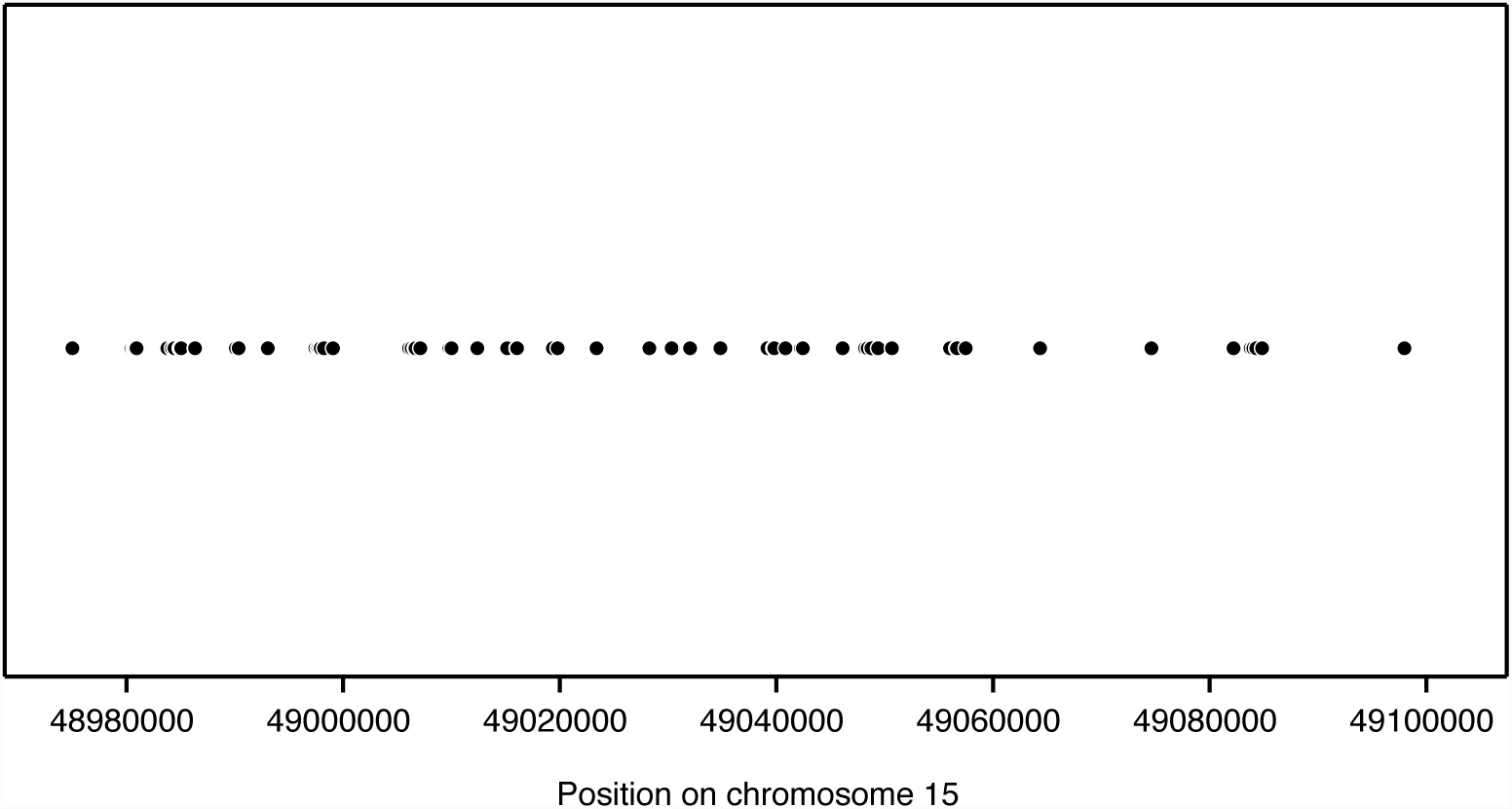
**Mapping locations in the *B. taurus* genome of reads used to seed *O. virginianus* HBB_A_ local assembly.**

**Fig. S8.**
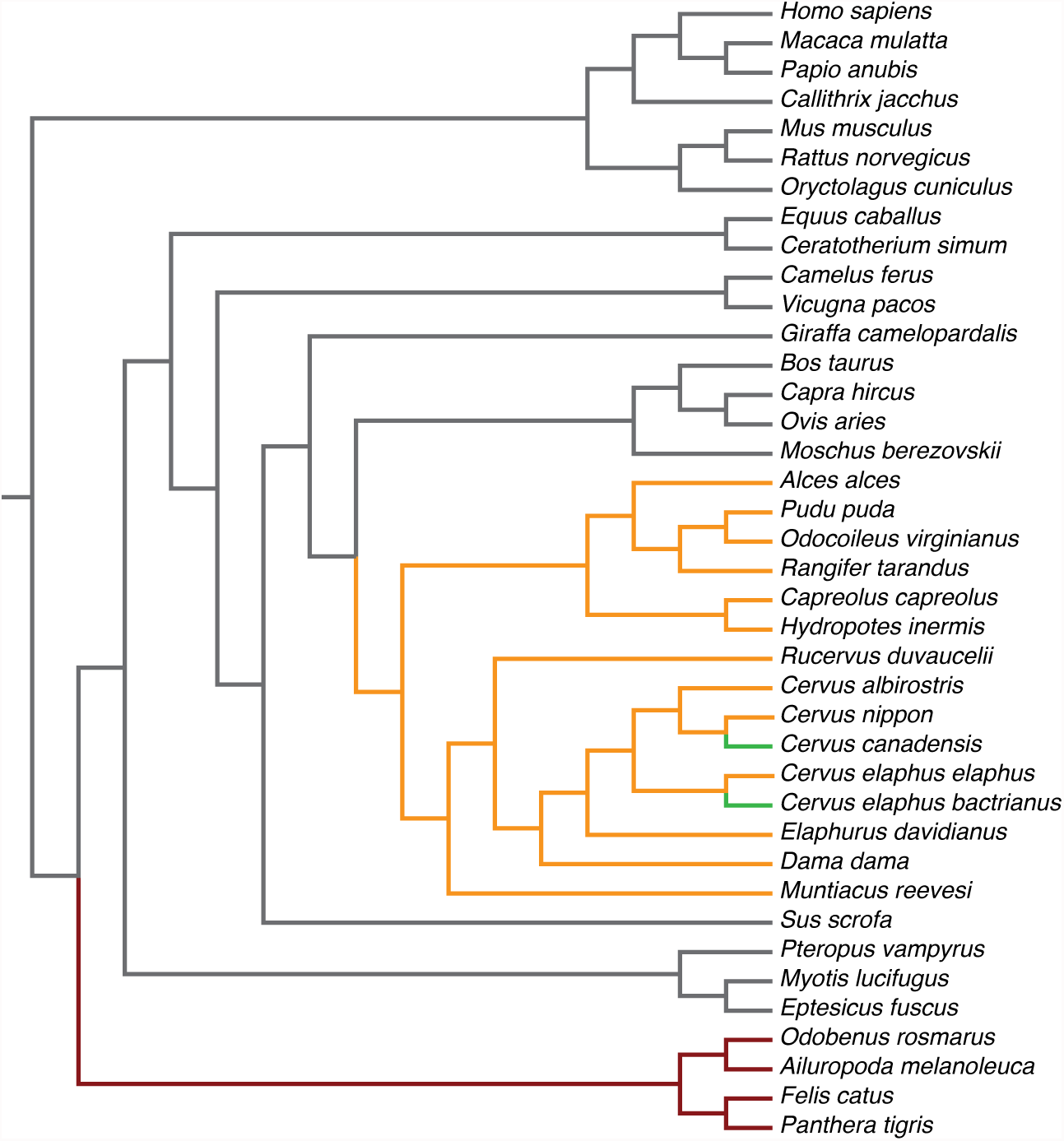
Cladogram of the mammalian species phylogeny used in this study. Coloured branches indicate deviations from and additions to the Timetree of Life phylogeny (see Materials and Methods); red: re-grafted Carnivora; orange: 10kTrees deer phylogeny; green: manually added branches absent from the 10kTrees phylogeny.

**Table S1.** Species considered in this study, previous evidence for sickling and sample origins. For each sample, species identity was confirmed by sequencing the mitochondrial *CytB* gene (see Table S2). For species in which both sickling and non-sickling individuals have been previously identified, the more common phenotype (as found in the associated references) is listed.

| Species | Common Name | Sample | Sample type | Source | Sickling state | References |
| --- | --- | --- | --- | --- | --- | --- |
| *Odocoileus virginianus* | White-tailed deer | WTD | Blood* | Penn State Deer Research Centre, Pennsylvania PA 16802, USA | Sickling† | 8,10,15,18,20,24,75 |
| *Dama dama* | Fallow deer | BFD | Blood* | ZSL Whipsnade Zoo, Whipsnade, Dunstable, LU6 2LF, UK (ZSL) | Sickling | 9-11,38 |
| *Rucervus duvaucelii* | Swamp deer | SD | Blood* | ZSL | Sickling† | 11,38,75 |
| *Elaphurus davidianus* | Père David's deer | PDD | Blood* | ZSL | Sickling | 9-11,38 |
| *Cervus nippon* | Sika deer | SIKA | Blood* | ZSL | Sickling† | 9-11,23,38,75,76 |
| *Cervus elaphus elaphus* | Red deer | RED | Blood | RZSS Highland Wildlife Park, Kincraig, Kingussie, PH21 1NL, UK (RZSS) | Sickling | 9-11,17,38 |
| *Cervus elaphus bactrianus* | Bactrian deer | BACT | Blood | RZSS | Indeterminate |  |
| *Cervus albirostris* | Whitelipped deer | WLD | Blood | RZSS | Indeterminate |  |
| *Alces alces* | European elk (Moose) | ELK | Blood / Muscle tissue | RZSS / Kezie Foods, Duns, TD11 3TT, UK | Does not sickle | 9,11,38 |
| *Rangifer tarandus* | Reindeer | REIN | Blood / Muscle tissue | RZSS / Kezie Foods, Duns, TD11 3TT, UK | Does not sickle | 9-11,38 |
| *Muntiacus reevesi* | Reeve's muntjac | MUNT | Muscle tissue | The Wild Meat Company, Woodbridge, IP12 2DY, UK | Sickling | 8,11,38 |
| *Pudu puda* | Pudu | PUDU | Blood | Bristol Zoo, Bristol Zoo Gardens, Clifton, Bristol, BS8 3HA, UK | Sickling | 38,74 |
| *Capreolus capreolus* | Roe deer | ROE | Tissue | V. Savolainen | Indeterminate |  |
| *Hydropotes inermis* | Chinese water deer | CWD | Tissue | V. Savolainen | Indeterminate‡ | 10,38 |
| *Cervus canadensis* | Wapiti (Elk) | WAP | Genomic DNA | East Stroudsburg University, 200 Prospect St, East Stroudsburg, PA 18301, USA | Does not sickle† | 9-12,38 |

\*Fresh blood samples processed with the PAXgene Blood DNA kit.
†Both sickling and non-sickling adult individuals previously recorded.
‡Only one individual tested (non-sickling). Conservatively listed as indeterminate.

